# Sex differences and effects of estrous stage on hippocampal-prefrontal theta communications

**DOI:** 10.1101/2020.05.16.099739

**Authors:** Kristofer J. Schoepfer, Yiqi Xu, Aaron A. Wilber, Wei Wu, Mohamed Kabbaj

## Abstract

Effective communication between the mammalian hippocampus and neocortex is essential to certain cognitive-behavioral tasks critical to survival in a changing environment. Notably, functional synchrony between local field potentials (LFPs) of the ventral hippocampus (vHPC) and the medial prefrontal cortex (mPFC) within the theta band (4-12 Hz) underlies innate avoidance behavior during approach-avoidance conflict tasks in male rodents. However, the physiology of vHPC-mPFC communications in females remains unestablished. Furthermore, little is known about how mPFC subdivisions functionally interact in the theta band with hippocampal subdivisions in both sexes in the absence of task demands. Given the established roles of biological sex and gonadal hormone status on innate avoidance behaviors and neuronal excitability, here, we characterize the effects of biological sex and female estrous stage on hippocampal-prefrontal theta signaling in freely-moving female and male rats. LFPs from vHPC, dorsal hippocampus (dHPC), mPFC-prelimbic (PrL), and mPFC-infralimbic (IL) were simultaneously recorded during spontaneous exploration of a familiar arena. Data suggest that theta phase and power in vHPC preferentially synchronize with PrL; conversely, dHPC and IL preferentially synchronize. Males displayed greater vHPC-PrL theta synchrony than females, despite similar regional frequency band power and inter-regional coherence. Additionally, several significant estrous-linked changes in hippocampal-prefrontal theta dynamics were observed. These findings support the hypothesis that biological sex and female estrous stage can both affect hippocampal-prefrontal theta signaling in a familiar environment. These findings establish novel research avenues concerning sexual dimorphisms and effects of gonadal hormone status in HPC-mPFC network activity pertaining to threat evaluation biomarkers.

**SIGNIFICANCE STATEMENT:** Effective signaling between subregions of the hippocampus and the prefrontal cortex underlies several cognitive-behavioral processes necessary for survival. Theta-frequency synchrony between the local fields of the ventral hippocampus and the prefrontal cortex regulates innate avoidance behavior in male rodents. However, this circuit remains understudied in females, despite its potential utility in research modeling the neurobiology of anxiety disorders (and sex differences therein). Here, we demonstrate that both biological sex and female hormone status have multifaceted effects on hippocampal-prefrontal theta-frequency signaling in freely-moving rats during exploration of a familiar arena. These findings provide insight into how biological sex and female estrous stage shape the way in which these circuits may represent the potential for threat.

## INTRODUCTION

The brain’s ability to adaptively respond to the body’s surroundings relies on synchronous oscillatory activity in the local field potentials (LFPs) of interconnected brain regions (1). In mammals, LFP interactions between subregions of the hippocampus (HPC) and the medial prefrontal cortex (mPFC) play a critical role in many cognitive functions critical for survival, for example, encoding and retrieving environmental cue information associated with a goal-directed behavior (2, 3). Indeed, disrupted HPC-mPFC interactions have been linked with several psychiatric disease states and may contribute to their pathophysiology (4). Along the dorsal-ventral (septal-temporal) axis of the hippocampus, multiple overlapping gradients of gene expression and anatomical connectivity are believed to produce distinct functional processes and behavioral outputs (5, 6). In rats, the ventral hippocampus CA1 (vHPC) sends bilateral, long-range, glutamatergic projections directly to ipsilateral dorsal subregions of the mPFC: namely, the prelimbic cortex (PrL) and infralimbic cortex (IL) (7–9). Similarly, in humans, the hippocampus is connected to regions of the medial prefrontal cortex by white matter tracts(10). Unlike the vHPC, neurons in the rat dorsal hippocampus CA1 (dHPC) do not directly project to the mPFC, instead primarily relaying first in midline nuclei of the thalamus (11).

Synchronized functional connectivity in the vHPC-mPFC circuit is believed to underlie the processing of environmental and subjective cues in order to assign emotional valence to the environment and guide an appropriate behavioral response based on perceived threat (12). Several findings in male mice have demonstrated that theta frequency (~8 Hz) phase synchrony in vHPC-mPFC LFPs underlies innate avoidance behavior in approach-avoidance conflict tasks such as the elevated plus maze (13–15). These findings have direct relevance to understanding the neurobiology of innate threat processing, and are of value in modeling “anxiety-like” states in rodents, as homologous brain regions to the rodent HPC and mPFC are involved in human expression of anxiety states and recall of extinguished fear (16–19).

Anxiety is a naturally-occurring adaptive response to a potential threat. However, when overexpressed in inappropriate contexts, it can result in an anxiety disorder: a class of psychiatric disease that disproportionately affects women (20). Sex differences in mammalian behaviors are due, in part, to the developmental and activational effects of circulating gonadal steroid hormones on the structure, and therefore the function, of neurons and other cells in the central nervous system (21). Along with other interconnected limbic structures including the amygdala, neurons in the HPC and mPFC are rich with receptors for androgens, estrogens, and progesterone (22, 23). Furthermore, natural cycling of estradiol and progesterone dynamically alters innate avoidance (“anxiety-like”) behaviors in female rodents (24–26) as well as functional connectivity between the HPC and mPFC in women (27). Together, this suggests a potential role for the hormonal milieu to act upon vHPC-mPFC circuits or networks in rodents to modify emotional reactivity to potential threat and alter the expression of innate avoidance behaviors. Surprisingly, however, the degree to which biological sex and/or female hormonal status may affect the coherent activity within HPC-mPFC networks has not yet been identified.

Here, we describe the roles of biological sex and female estrous stage on the neural oscillations within and between the mPFC and HPC of awake freely-moving rats during active exploration of a familiar arena. Also, we compare functional connectivity patterns between subregions of the mPFC (PrL and IL) and subregions of the hippocampus (dorsal and ventral CA1).

## RESULTS

Adult male and female rats were chronically implanted with electrode bundles targeting mPFC-infralimbic (IL), mPFC-prelimbic (PrL), ventral hippocampus CA1 (vHPC), and dorsal hippocampus CA1 (dHPC) (**Fig. 1a**). Male and female implant locations were matched (**Suppl. Fig. S1**). After recovery, LFPs from the four brain regions were simultaneously recorded daily for 10-minute trials in a familiar arena during freely-moving exploratory behavior (**Fig. 1b**).

**FIGURE 1.**
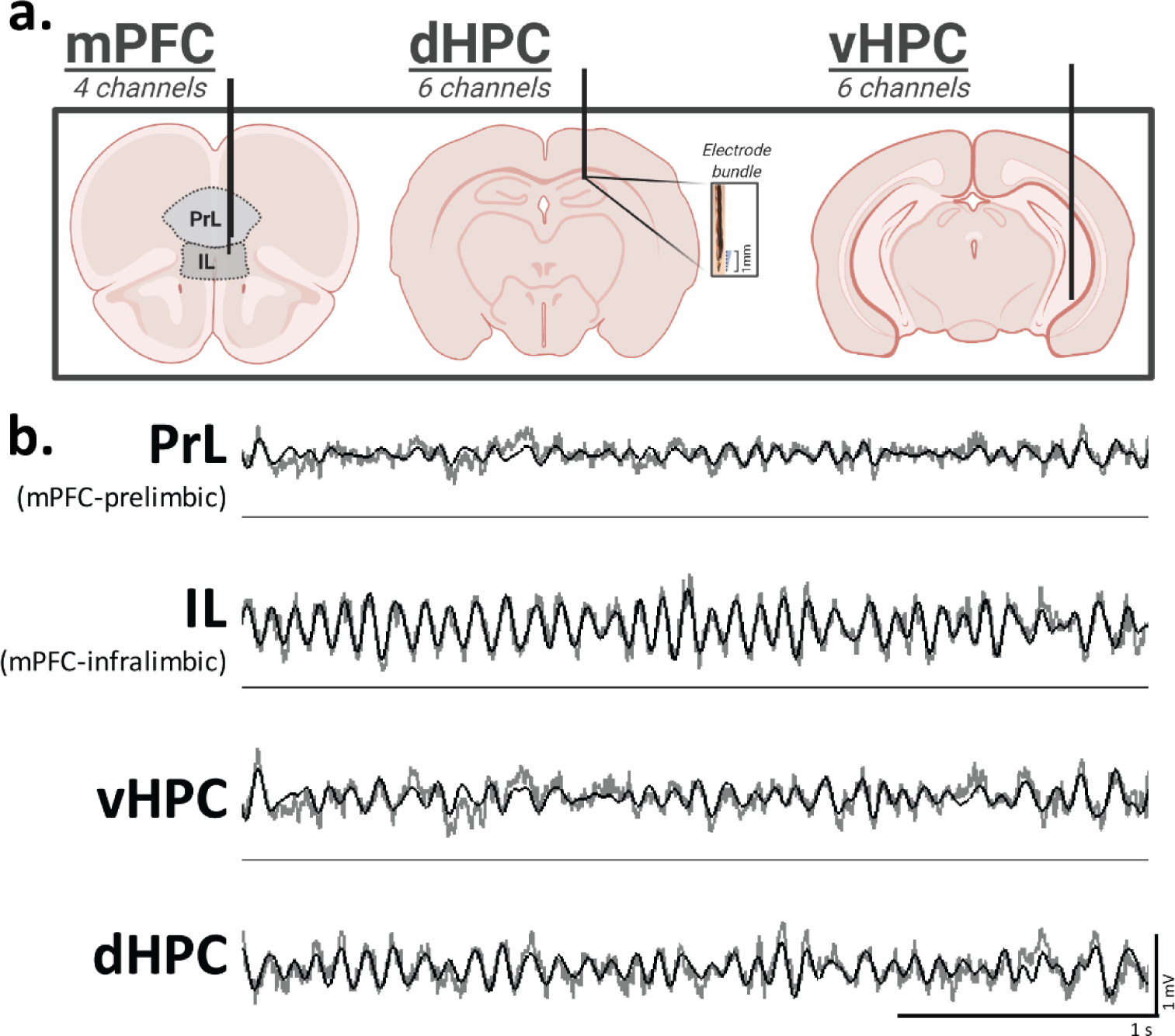
Electrode implant sites and representative LFP traces. **(a)** Illustration of multi-site intracranial electrode bundle implants. Bundles of six wires with a vertical spread of 1mm were targeted to vHPC CA1 and dHPC CA1. Bundles of four wires were targeted to IL and PrL. Male and female implant locations are matched, **(b)** Representative traces of simultaneously-recorded LFPs from IL, PrL, dHPC, and vHPC during active exploration in a familiar arena. Raw traces are plotted in gray and theta-filtered traces are overlaid in black.

### No differences in regional power spectra between sexes or estrous stages

For each subject, LFP data from all epochs of 5-15 cm/s movement in the familiar arena were concatenated. Power spectral density (PSD) estimates were calculated from simultaneous PrL, IL, vHPC, and dHPC LFPs in the familiar arena. For each brain region, the function (shape) of broadband (0.5-100 Hz) PSDs were analyzed as a function of biological sex, then female data were analyzed separately as a function of estrous stage. There were no effects of biological sex or female estrous stage in the function of 0.5-100 Hz PSD estimates in all four brain regions (**Fig. 2a-d**). Similarly, when the PSD were separated into biologically-relevant frequency bands (delta 1-4 Hz, theta 4-12 Hz, beta 15-30 Hz, gamma 30-80 Hz), no effects of biological sex or estrous stage were observed in any brain region (ps>0.05).

**FIGURE 2.**
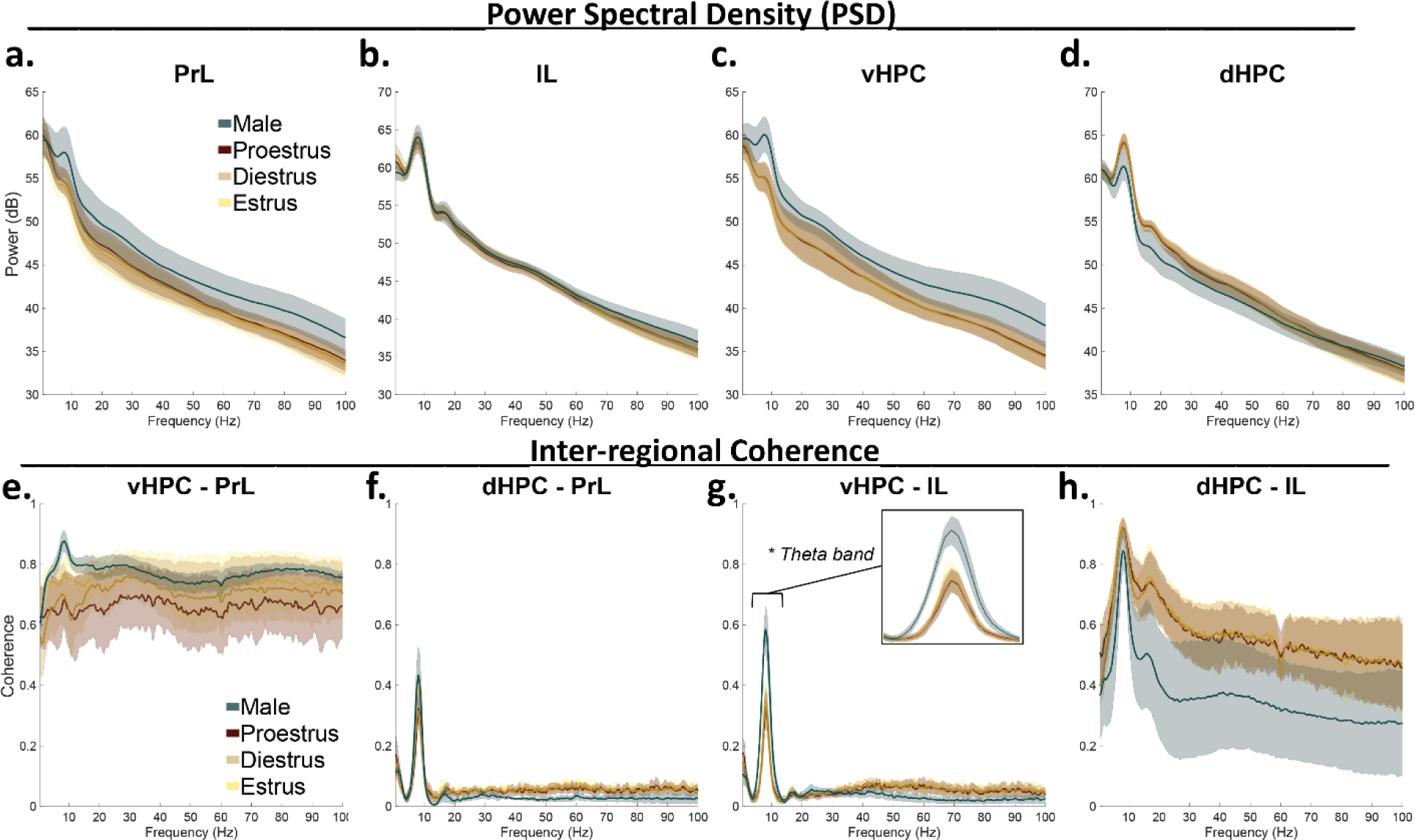
No differences in PSDs, but a sex-associated difference in coherence. **(a-d)** Power spectral density estimates in prefrontal and hippocampal subregions. Neither biological sex nor estrous stage significantly affected the shape of PSD estimates (0.5-100 Hz) in **(a)** PrL (sex p=0.2745; estrous p=0.5600; male vs estrous p=0.3525), **(b)** IL (sex p=0.7354; estrous p=0.9700; male vs estrous p=0.9676), **(c)** vHPC (sex p=0.1096; estrous p=0.9778; male vs estrous p=0.0517), or **(d)** dHPC (sex p=0.2031; estrous p=0.9995; male vs estrous p=0.5359). **(e-h)** Coherence between hippocampal-prefrontal pairs. Neither biological sex nor estrous stage significantly affected the shape of magnitude-squared coherence (0.5-100 Hz) between **(e)** PrL and vHPC (sex p=0.5744; estrous p=0.8990; male vs estrous p=0.8912), **(f)** PrL and dHPC (sex p=0.2688; estrous p=0.9444j male vs estrous p=0.6023)7 or(h) IL and dHPC (sex p=0,3194; estrous p=0.9971; male vs estrous p=0.55GB). **(g)** Coherence between vHPC and IL (0.5-100 Hz) was not affected by estrous stage (p=0.9908), but differed significantly between the sexes (*p=0.0209), an effect driven by the theta band (4-12 Hz, *p=0.0l82; shown in inset). Data represent epochs of movement 5-15 cm/s and are plotted as mean + SEM; shaded error represents subjects. Female-male comparisons were analyzed by functional two-sample F-tests, with significant p-values *p<0.05, Effects of estrous stage were analyzed by functional ANOVAs. Significant p-values in estrous stage comparisons were *p<0.0167, adjusted to account for 3 pairwise comparisons (proestrus/diestrus, estrus/diestrus, estrus/proestrus, any stage versus males). Coherence was analyzed between 0.5-100 Hz, then frequency bands of interest were analyzed separately. Sample sizes are found in Table 1.

**Table 1.** Sample sizes. Biological and technical replicates included in this study.

|  | FEMALE |  |  |  |  | MALE |  |  |  |
| --- | --- | --- | --- | --- | --- | --- | --- | --- | --- |
| *Subjects:* | "F1" | "F2" | "F3" | "F4" | "F5" | "M1" | "M2" | "M3" | "M4" |
| *n(Trials):* | 25 | 28 | 23 | 26 | 28 | 27 | 27 | 27 | 25 |
| *n(Trials.Diestrus):* | 13 | 14 | 14 | 13 | 12 |  |  |  |  |
| *n(Trials.Proestrus):* | 3 | 9 | 3 | 7 | 6 |  |  |  |  |
| *n(Trials.Estrus):* | 6 | 2 | 5 | 5 | 9 |  |  |  |  |
| *n(Trials.Metestrus):* | 3 | 3 | 1 | 1 | 1 |  |  |  |  |

### Sex-associated difference in inter-regional coherence without estrous modulation

Coherence in frequency-specific LFPs between brain regions has been theorized to play a fundamental role in coordinating local neural computations (28). Coherence between hippocampal-prefrontal LFP pairs was calculated from rats moving 5-15 cm/s in a familiar arena; all epochs were concatenated by subject. There was a significant sex-associated difference in the function of 0.5-100 Hz coherence between vHPC and IL, an effect driven by increased coherence for males within the theta band (**Fig. 2g**). No other hippocampal-prefrontal pairs differed in 0.5-100 Hz coherence between the sexes, nor within the theta, delta, beta, or gamma frequency bands (**Fig. 2e,f,h**). There were no effects of estrous stage on the function of hippocampal-prefrontal coherence in any frequency range.

### Hippocampal-prefrontal theta band power correlations differ by sex and estrous stage

To determine how instantaneous theta band amplitudes synchronize hippocampal and prefrontal subregions, theta band power correlations between hippocampal-prefrontal pairs were calculated over time, and the correlation coefficients were collected per trial (**Fig. 3a**). When comparing subject means, theta band power correlations were significantly enhanced in vHPC-PrL and dHPC-IL circuits as compared to dHPC-PrL and vHPC-IL circuits (**Fig. 3b**). Correlation coefficients were then analyzed as a function of biological sex and as a function of the female estrous stage using a hierarchical bootstrapping method of data resampling(**29**). Theta band power correlations between vHPC-PrL were significantly greater in males than females, but female estrous stage had no significant impact (**Fig. 3c**). Theta band power correlations between dHPC-PrL were not significantly affected by biological sex or female estrous stage (**Fig. 3d**). However, proestrus females showed a reduction in dHPC-PrL theta power correlations as compared to diestrus females and males that approaches statistical significance. Similarly to the vHPC-PrL circuit, vHPC-IL theta band power correlations were significantly greater in males as compared to females (**Fig. 3e**), suggesting theta band signaling in vHPC-mPFC circuits is of greater amplitude in male rats as compared to females. Females in estrus displayed a significant decrease in vHPC-IL theta power correlations as compared to diestrus females, with proestrus values ranging between them. Lastly, dHPC-IL theta band power correlations were significantly greater for females as compared to males (**Fig. 3f**). Proestrus stage significantly reduced dHPC-IL theta power correlations in females but remained distinguishable from males. These data suggest that in females, proestrus stage downregulates dHPC-mPFC theta power correlations, while these signals remain constant throughout diestrus and estrus.

**FIGURE 3.**
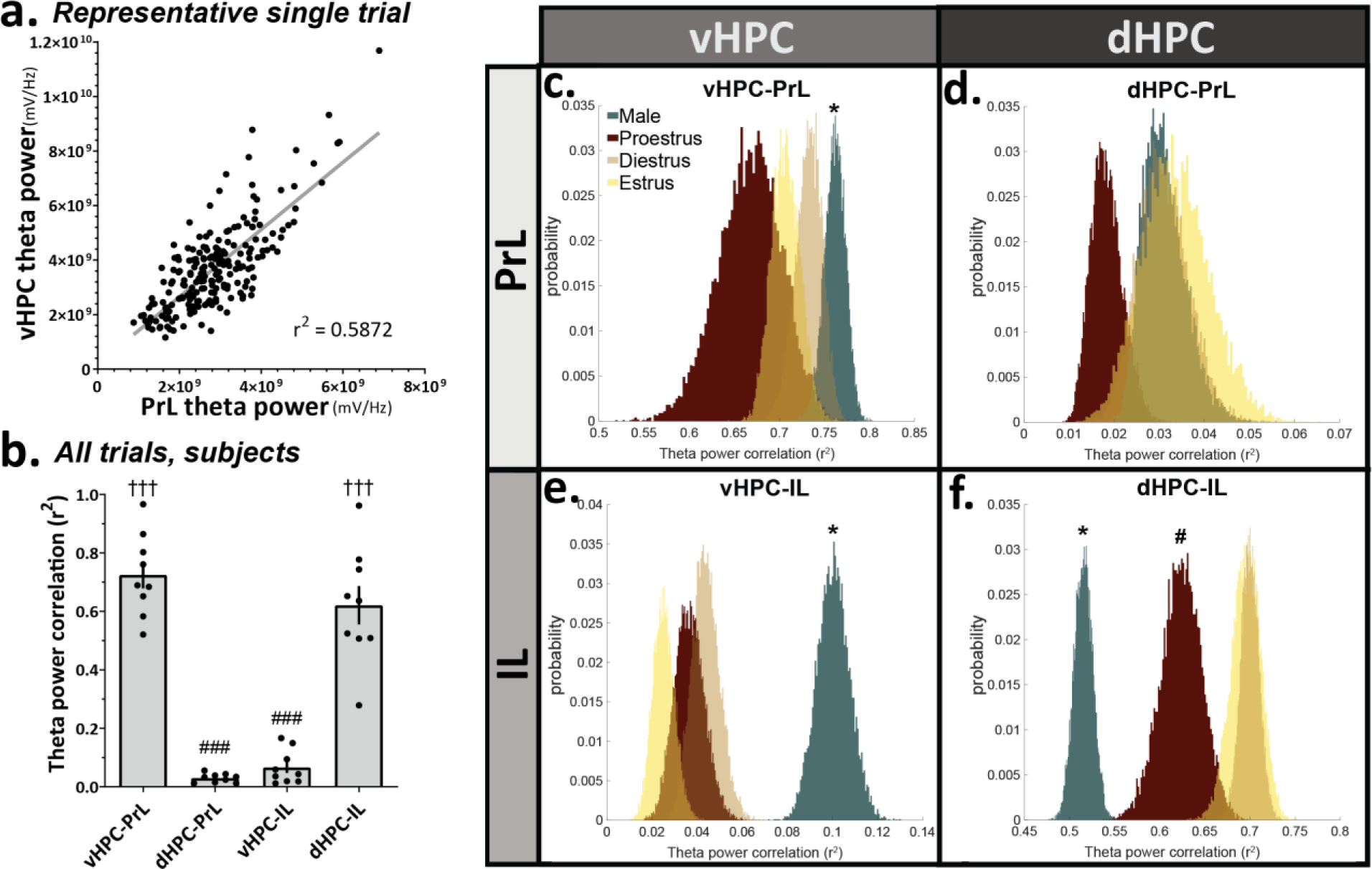
Hippocampal-prefrontal theta band power correlations differ by sex and estrous stage. **(a)** Representative example of one trial’s theta power correlation between vHPC and PrL in the familiar arena. Each data point represents the sum of theta band power in a 2.6sec window, **(b)** Subject-averaged theta power correla¬tions between vHPC-PrL and dHPC-IL are significantly greater than those between dHPC-PrL or vHPC-IL (RM 1-way ANOVA ***p<0.0001; Tukey’s multiple comparison test, ###p<0.0001 vs vHPC-PrL, †††p<0.0001 vs dHPC-PrL; n = 9 subjects), **(c)** vHPC-PrL theta power correlations were significantly greater for males than females (*p=0.0022). Female estrous stage had no significant effect on vHPC-PrL theta power correlations (diestrus/proestrus p=0.0768; diestrus/estrus p=0.2564; proestrus/estrus p=0.2737). **(d)** dHPC-PrL theta power correlations were not significantly affected by biological sex (p=0.6584) or by female estrous stage (diestrus/proestrus p=0.0515; diestrus/estrus p=0.7490; proestrus/estrus p=0.0476). **(e)** vHPC-IL theta power correlations were significantly greater for males than Females (*p<l-10), and this was also true for each female estrous stage (males vs diestrus *p<l-4; vs proestrus *p<l^−4^; vs estrus *p<l^−5^). For females, estrus significant¬ly decreased vHPC-IL theta power correlations as compared to diestrus stage (*p=0.0147), but no other significant differences were observed (diestrus/proestrus p=0.4362; proestrus/estrus p=0.1702). **(f)** dHPC-IL theta power correlations were significantly greater for females than males (*p<l^−10^), and this was also true for each female estrous stage (males vs diestrus *p<l^−10^; vs proestrus *p<l^−7^; vs estrus *p<l^−10^). While female diestrus and estrus stages were comparable (p=0.8409), proestrus significantly reduced dHPC-ILtheta power correlations in females (diestrus/proestrus #p=0.0012; proestrus/estrus *p=0.0081). Data were analyzed via direct probabili¬ty estimates on hierarchically-bootstrapped samples; p_boot_ values were converted to two-tailed p-values. Significant p-values in male-female comparisons are *p<0.05. Significant p-values in estrous stage comparisons are *p<0.0167, adjusted to account for 3 pairwise comparisons (proestrus/diestrus, estrus/diestrus, estrus/proestrus, any stage versus males). Sample size inputs for hierarchical bootstrapping are found in Table 1.

### Hippocampal-prefrontal theta phase lags differ by sex and estrous stage

Theta phase synchrony was determined by computing the distributions of theta band phase lag differences between prefrontal-hippocampal pairs, then comparing the bootstrap-sampled widths at ½ the maximum distribution peaks. Calculated this way, small values indicate a sharp peak with reliable phase lags between hippocampal-prefrontal pairs, whereas large values indicate unreliable theta phase lag distributions between brain regions (**Fig. 4a**). Combining all trials and subjects, theta phase lags from vHPC to PrL and from dHPC to IL were more reliably distributed (smaller width at half-max peak values) than other connections, suggesting that theta phase synchrony may represent a preferential mode of communication in these circuits (vHPC-PL and dHPC-IL) compared to the other circuits assessed (vHPC-IL and dHPC-PrL; **Fig. 4b**). In the vHPC and PrL circuit, theta phase lags were also more tightly distributed in males as compared to females (**Fig. 4c**). Females in diestrus and proestrus were similar, but estrus significantly enhanced theta phase synchrony between vHPC-PrL versus proestrus, such that estrus females were not significantly different from male values. Conversely, theta phase lags in the dHPC to PrL circuit were more tightly distributed in females than in males (**Fig. 4d**). In females, estrus stage also significantly improved dHPC-PrL theta phase synchrony as compared to diestrus stage but was not different from proestrus. Between vHPC and IL, theta phase lags were more tightly distributed in males as compared to females (**Fig. 4e**). Females in diestrus and estrus were similar, but proestrus significantly reduced vHPC-IL theta phase lag consistency as compared to all other groups. Lastly, theta phase lag distributions from dHPC to IL were significantly different between all groups, such that diestrus females had the most reliable theta phase synchrony, followed by proestrus females, estrus females, then males (**Fig. 4f**). Overall, females’ dHPC-IL theta phase lags were more reliably distributed than males’.

**FIGURE 4.**
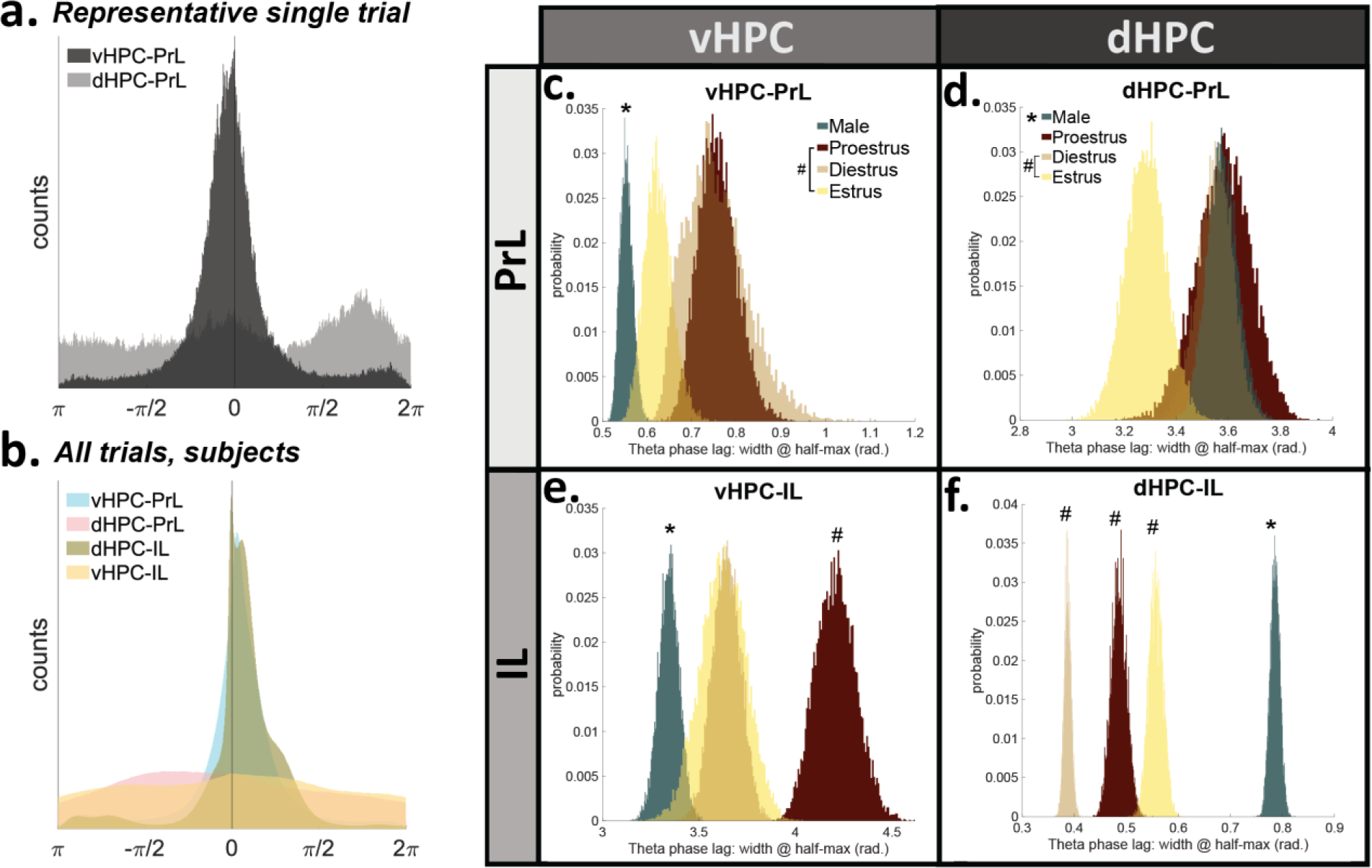
Hippocampal-prefrontal theta phase lags differ by sex and estrous stage. **(a)** Representative distributions of hippocampal-prefrontal theta phase lags from one trial: reliable theta phase synchrony from vHPC to PrL, but unreliable phase synchrony from dHPC to PrL. **(b)** Combining all trials, theta phase lags from vHPC to PrL (0.6193 rad.) and dHPC to IL (0.5194 rad.) were more reliable than those from dHPC to PrL (5.7238 rad.) and vHPC to IL (6.2632 rad.), **(c)** Males had greater theta phase synchrony (i.e. smaller width at half-max distribution peak) than females between vHPC and PrL (*p<l^−10^). Estrous stage dynamically modulated vHPC-PrL theta phase synchrony, such that estrus females were significantly different from proestrus females (*p=0.0102), but were not significantly different from diestrus females or males (estrus/diestrus p=0.0539; estrus/males p=0.0204; diestrus/proestrus p=0.9095). **(d)** Females had greater phase synchrony than males from dHPC to PrL (*p=0.0005). Female estrus stage trials had significantly enhanced dHPC-PrL theta phase synchrony as compared to diestrus female trials (*p=0.0106) or male trials (*p=0.0022), but were not significantly different from proestrus female trials (p=0.0262). **(e)** Males had greater phase synchrony than females from vHPC to IL (*p<l^−10^). While female diestrus and estrus stages were comparable (p=0.9305), proestrus significantly reduced vHPC-IL theta phase synchrony versus all other groups (vs diestrus *p<l^−7^; vs estrus *p=0.0102; vs males *p<l^−10^). **(f)** Females had greater theta phase synchrony than males from dHPC to IL (*p<l^−10^), and each female hormone stage was significantly different from males (*ps<l^−10^). Estrous stage significantly modulated dHPC-IL theta synchrony in females in an order of Diestrus > Proestrus < Estrus (diestrus/proestrus *p<l^−10^; diestrus/estrus *p<l^−10^; proestrus/estrus *p<l^−10^). Data were analyzed via direct probability estimates on hierarchical¬ly-bootstrapped samples; p_boot_ values were converted to two-tailed p-values. Significant p-values in male-female comparisons are *p<0.05. Significant p-values in estrous stage comparisons are *p<0.0167, adjusted to account for 3 pairwise comparisons (proestrus/diestrus, estrus/diestrus, estrus/proestrus, any stage versus males). Sample size inputs for hierarchical bootstrapping are found in Table 1.

## DISCUSSION

Adult male and female rats were placed into a familiar arena, and LFPs from PrL, IL, vHPC, and dHPC were simultaneously recorded over many trials. Within the areas studied, neither biological sex nor female estrous stage significantly affected the power across a range of known brain rhythms. While plasticity in the vHPC-mPFC pathway has been documented (30), the female estrous stage had no effect on coherence between any hippocampal-prefrontal pairs in our study, suggesting natural fluctuations in estradiol and progesterone alone may be insufficient to elicit observable plasticity during exploratory behaviors. Interestingly, males had greater theta band coherence than females between vHPC and IL (**Fig. 2g**). In fact, our data suggest that males had overall greater effective theta signaling between vHPC and both mPFC subregions versus females. Males had the greatest theta power correlations between vHPC and IL/PrL (**Figs. 3c,3e**), as well as the most reliable theta phase lag distributions between vHPC and IL/PrL (**Figs. 4c,4e**). These findings align with the vHPC’s known monosynaptic projections to mPFC in the male rat, whereas dHPC-mPFC connections are believed to be polysynaptic (**9**). Conversely, females had greater effective theta signaling between the dHPC and mPFC subregions, though this effect appears to be stronger for IL than PrL. It is worth noting here that the referenced neuroanatomy studies utilized male rat subjects, and literature examining the anatomy of HPC-mPFC monosynaptic projections in females is extremely limited. Several possible explanations for the observed male-female differences in vHPC-mPFC theta signaling exist. First, fewer excitatory vHPC neurons may project monosynaptically to IL and PrL in females as compared to males. Second, males may have greater vHPC theta power than females, which would drive synchronization of vHPC-mPFC communications differentially. However, no sex differences in vHPC power were found here (**Fig. 2c**). Third, males could have enhanced anatomical and/or synaptic connectivity with other corticolimbic brain regions that resonate in theta with the mPFC and vHPC (for example, the basolateral amygdala). Lastly, there could be sex-differential hippocampal connectivity with the medial septum or with the fimbria, believed to pace theta rhythms in male rats within dHPC and vHPC, respectively. However, additional anatomical studies are needed to fully explain the observed sex-associated differences in vHPC-mPFC theta connectivity.

To our knowledge, these data are the first to demonstrate several significant changes in HPC-mPFC theta signaling across the female estrous cycle in rats. During diestrus, when circulating levels of progesterone and estradiol are low, IL theta phase preferentially synchronizes with dHPC theta phase and PL disengages, while vHPC-IL theta power correlations are at their strongest point. During the subsequent proestrus stage, when progesterone and estradiol concentrations surge, dHPC-IL theta phase synchrony increases at the expense of vHPC-IL phase synchrony, and dHPC power correlations with mPFC are reduced. Finally, during estrus stage, when circulating estradiol remains elevated but progesterone levels are diminished, PrL hits peak phase synchronization with HPC, and vHPC-IL power correlations increase without modifying phase synchrony. For female rats, there is likely an evolutionary benefit to having a dynamic corticolimbic system that adaptively changes the strengths of effective signaling along with the “tides” of endogenous hormone cycles. Notably, proestrus stage signals sexual receptivity in female rats (21), aligning with reported increases in open-zone exploration during approach-avoidance conflict tasks (25, 26, 31). Therefore, one explanation may be that the hormonal surge during proestrus shapes communications within corticolimbic circuits to reduce threat bias, thereby promoting exploratory behavior and improving the odds of successfully mating. However, it is important to emphasize that natural selection acts on behaviors essential for survival, rather than the biological mechanisms responsible for generating those behaviors (32), so future research including task engagement is justified.

Another possible explanation for the observed differences in HPC-mPFC theta connectivity is that receptors for estrogen, progesterone, and androgens are expressed differentially between the sexes and across the female estrous stage within corticolimbic structures (22, 23, 33–35). Therefore, the developmental and/or activational effects of gonadal hormones (e.g. estradiol, progesterone, and testosterone) on the HPC, mPFC, and/or interconnected limbic structures may play a role in sex- and estrous-related changes in theta signaling.

The vHPC-PrL circuit has been studied in the context of approach-avoidance behaviors to better understand its utility for identifying potential biomarkers of pro-avoidant states, ie, modeling anxiety-like states in rodents. Previous studies suggest that in male mice, vHPC-PrL theta phase synchrony may serve as a functional biomarker for innate avoidance behavior in approach-avoidance conflict tasks such as the elevated plus maze (13–15). Extrapolating this concept, one might hypothesize that male rodents would have overall greater vHPC-PrL theta band coupling than females, given that male rodents spend more time than females in the closed portions of the elevated plus maze when the estrous stage is ignored (36). Our findings support the hypothesis that males have overall greater vHPC-PrL theta band coupling than females in measurements of power correlation (**Fig. 3c**) and phase synchronization (**Fig. 4c**) during exploration of a familiar arena. However, variability between subjects was too large to observe a significant effect of sex in vHPC-PrL coherence (**Fig. 2e**). Along with its documented role in innate avoidance behaviors, the vHPC-PrL circuit has recently been linked with fear suppression via learned safety cues (37), suggesting a potential extension of these findings to learned avoidance behaviors. However, additional studies are needed to determine the role of vHPC-PrL signaling in females in this context.

Together, data suggest that while maintaining similar power spectra within each brain region, biological sex and female estrous stage selectively modify synchronized theta frequency communications between hippocampal and prefrontal subregions during active exploration of a familiar arena. We found that theta signaling is most cohesive in vHPC-PrL and dHPC-IL circuits overall. In male rats, vHPC-mPFC theta signaling was more tightly coupled than dHPC-mPFC signaling, whereas the opposite was true for females, most notably in IL. These findings show that even in the absence of task demand, biological sex and the female estrous stage can both affect synchronized theta communications between the hippocampus and prefrontal cortex in-vivo, with potential implications for biomarkers of threat representation.

## METHODS

### Animal subjects

Adult male (n = 4) and female (n = 5) Sprague-Dawley rats aged eight weeks obtained from Charles River Laboratories (Raleigh, NC) were used in this study. Animals were housed in same-sex pairs with environmental enrichment for 3+ days before surgery and were singly-housed with environmental enrichment after surgery. Animals were kept under standard controlled temperatures, reversed 12-hr light-dark cycles (lights off at 10:00 am), and were allowed ad libitum access to standard chow and water. All procedures were carried out under strict accordance with the NIH Guide for the Care and Use of Laboratory Animals, and the animal protocol was approved by the Florida State University Institutional Animal Care and Use Committee.

### Electrode construction

Formvar-insulated nickel-chromium alloy wire (California Fine Wire, 0.003 in) was used to construct custom electrode probes. Six-electrode bundles were constructed by twisting wire and adhering with a heat gun. The brain-oriented bundle end was cut with a fresh scalpel at an angle such that the uninsulated tips of the 6 wires covered a 1mm spread. Four-electrode bundles were constructed similarly but with 1mm spacing between each wire (total spread = 3mm), and adhering them with glue away from the tips. The free ends of all bundles were fire-stripped of insulation and were mechanically coupled to gold pin receptacles (Mill-Max, #0489-0-15-15-11-27-04-0) using Tygon ND-100-80 tubing (0.01 in ID; 0.03 in OD). A short piece of insulated wire serving as a reference electrode was coupled to a pin receptacle. Electrode impedances were tested in saline (nanoZ, White Matter, 1004 Hz, 40 cycles); only bundles with impedances of 0.1-0.5 MΩ were used. Channels were line-labeled in their bundle by passing current (−2 μA) into each wire in saline and mapping the resultant air bubble. An insulated micro coaxial ground wire (42 AWG, Molex #100065-0023) was soldered to a gold pin receptacle and a stainless-steel skull screw (Fine Science Tools, #19010-00).

### Surgery

Rats were deeply anesthetized with inhaled isoflurane and secured in a stereotaxic apparatus (Kopf Instruments). The skull angle was leveled (<50 μm) using bregma/lambda landmarks, 1-2 skull screws were manually driven into each bone plate, and the ground-coupled screw was implanted in the left frontal bone plate near the olfactory bulb. Craniotomies were made on the right hemisphere over the implant coordinates, and durotomies were performed. Electrode bundles were lowered into the brain with a micromanipulator targeting the following coordinates: mPFC: AP +3.0, ML +0.7, DV −4.0 (4 channels); dHPC CA1: AP −3.8, ML +2.5, DV-3.0 (6 channels); vHPC CA1: AP −5.5, ML +5.0, DV –6.6 (6 channels) (38). A reference wire was implanted in shallow cerebellar white matter (AP −12.08, ML +0.85, DV −0.3).

Anterior/posterior (AP) and medial/lateral (ML) coordinates are in reference to bregma, and dorsal/ventral (DV) coordinates are in reference to dura mater. Electrodes were affixed to the skull using C&B Metabond (Covetrus), then dental cement (Stoelting, #51458). The free ends of the electrodes were pinned out to a Mill-Max header cut to 9×2 size (Mill-Max, #853-93-100-10-001000), and all hardware was encased in dental cement, leaving the header ports open. Animals were monitored until emergence from anesthesia and were treated with analgesics and topical antibiotics as needed.

### Behavioral protocol

Animals were allowed to recover for at least 10 days or until regaining pre-surgery body weight and displaying a healthy scalp surrounding the implant. The recording arena consisted of a 46 x 46 x 55 cm gray acrylic box with an open top (Maze Engineers). Rats were acclimated to the recording arena, handling, and tethering to the headstage for three 10-min sessions daily for three days. Animals were transported in light-protected conditions to the dark recording room and were allowed to acclimate for 15 min. Staticide (ACL, Inc.) was applied to the arena’s floor and walls before each trial. LFPs from PrL, IL, vHPC, and dHPC were recorded for 10 min in the recording chamber (“familiar arena”). All recordings occurred between +3-5 hrs into the dark cycle and were performed in the dark. The arena was cleaned with 70% ethanol after each trial. Each animal provided 23-28 recordings (Table 1).

### Data acquisition

Local field potentials (LFPs) from freely-moving rats were recorded using a 16-channel unity-gain acquisition system (Neuralynx, DigiLynx 4SX) at 32 kHz sampling rate, referenced to the ground screw, and band-pass filtered 0.5-600 Hz. A monochrome video camera mounted overhead tracked an infrared LED mounted to the headstage preamplifier (XY coordinates) and captured video tracking data at 30 Hz.

### Estrous stage determination

Vaginal epithelial cells were collected daily via sterile saline lavage from female rats immediately after each LFP recording (~3hr into dark cycle). Samples were visualized under a 10X brightfield microscope; cell cytology informed estrous stage (39). To account for potential stress associated with the lavage technique, males were handled similarly daily.

### Data analysis

Data were imported into MATLAB (R2018b, Mathworks) and were analyzed with custom-written scripts, inbuilt functions, functions from the Communication Toolbox and Signal Processing Toolbox, and open-source packages. Data analysis scripts can be accessed on GitHub (https://github.com/KrisNeuro/KJS-Thesis). LFPs were high-pass filtered at 0.5 Hz, 60 Hz power line interference and 6 harmonics were attenuated (40), and LFPs were down-sampled to 2 kHz. Individual channels were excluded from further analysis if found to be poorly grounded or of significantly lower amplitude than other channels in that brain region across all trials. Of the remaining channels, for each trial, a matrix of phase-locking values (PLV) between 0.5-100 Hz was generated via the multitaper method (Chronux toolbox, http://chronux.org/ (41)) to compare all channels to each other. The mean PLV matrix was calculated, and channels within each brain region were visually inspected. Values ranged from 0-1; individual channels with PLV <0.6 versus all other channels in that brain region were considered outliers and were excluded from further analysis. In each brain region, the mean voltage of remaining usable channels was calculated at each time point, generating a precleaned “regional LFP”. XY coordinates from the headstage LED tracker were upsampled with padding to 2 kHz, co-registered with LFP timestamps, and converted to cm units. Animal linear movement velocities were calculated using the Freely Moving Animal Toolbox (http://fmatoolbox.sourceforge.net) smoothed with a Gaussian kernel (std. dev.= 1 sec). In freely-moving rodents, hippocampal theta activity increases with running speed (42). To reduce this variability, LFPs were velocity-filtered to exclude periods of movement slower than 5 cm/s or faster than 15 cm/s in analyses of power spectral density, frequency band power, and coherence. For each subject, epochs of movement 5-15 cm/s from all familiar arena recordings were concatenated into a single long time series containing four simultaneously-recorded pre-cleaned LFPs from PrL, IL, vHPC, and dHPC. Power spectral densities were calculated using the Welch method (moving window of 0.4s, 90% overlap between windows, 4000 nFFTs). Plotted power spectral data are decibel-transformed. For hippocampal-prefrontal coherence, 5-15 cm/s LFP data from each subject were concatenated, and magnitude-squared coherence was calculated for each circuit between 0.5-100 Hz, then separately for the theta band (4-12 Hz) (2s Hann window, 50% overlap). To calculate power correlations between areas, theta power was calculated for each brain region over time from the full trials, regardless of running velocity (multitaper cross-spectrogram, 2.5 sec window, no overlap between windows, NW=2.5, 2048 nFFTs). Then, two-tailed linear correlation coefficients (r^2^) were calculated from each hippocampal-prefrontal pair for each trial. To calculate theta phase lags, LFPs (all movement velocities included) were band-pass filtered for the theta band (4-12 Hz, Butterworth, MATLAB function ‘filtfilt’ to correct for phase) To ensure accurate theta phase, data included only epochs when dHPC theta power was greater than its mean for that recording. Hilbert transform was applied to the theta-filtered time series to extract instantaneous theta phase angles from each region. Values <–π or >π were adjusted to account for the oscillation wraparound. Instantaneous theta phase was subtracted between pairs of signals for each recording, and distributions of phase differences were analyzed, with values between -π and π in increments of π/100. Data were analyzed by calculating the width (in radians) of half the maximum phase lag distribution peak, such that tightly-coupled theta oscillations have a narrow peak width and poorly-coupled oscillations have a large peak width.

### Statistical analyses

Data were analyzed using MATLAB or Prism (8.1.3, GraphPad) software. Two-dimensional data (PSD, coherence) were analyzed using functional F-tests and functional ANOVAs to compare the function (shape) of the curves. Here, all trial epochs of 5-15 cm/s movement were concatenated by subject before calculating PSD or coherence, capturing all spectral data for that subject. For theta band power correlations and theta phase lags, comparisons between the sexes and between estrous stages were analyzed using a hierarchical bootstrapping approach adapted from Saravanan et al. (29). Trials were bootstrapped to subject, resampling data 10^4^ permutations with replacement. Bootstrapped samples were statistically compared via joint-probability matrix, directly assessing the probability of the second variable being equal to or greater than the first (p_boot_). One-tailed p_boot_ values were converted to two-tailed p-values via: 2*min(p_boot_, 1-p_boot_). To compare global differences in theta band power correlations and theta phase lags between hippocampal-prefrontal circuits, subject means were analyzed via repeated-measures 1-way ANOVA with Tukey’s post-hoc tests on normally-distributed data, or Friedman test with Dunn’s post-hoc tests on abnormally-distributed data. Alpha (p) was set to 0.05 when comparing independent data (males vs females). Significant p-values were set to 0.0167 in estrous stage comparisons to account for 3 pairwise comparisons (Proestrus/Diestrus, Estrus/Diestrus, Estrus/Proestrus; or any estrous group vs males). Metestrus trials were excluded in analyses of estrous stage effects due to small sample size. Statistical outputs and p-values are reported in the figure legends.

### Histology

After all recordings, animals were overdosed with sodium pentobarbital solution (Covetrus, Soccumb, delivered i.p.) and were transcardially perfused with paraformaldehyde (4% w/v in 100 mM phosphate-buffered saline, pH 7.4). Brain tissue was collected and post-fixed in 4% paraformaldehyde overnight at 4°C. Coronal brain tissue slices (40 μm) were collected using a vibratome (Leica VT1000S). Tissue was mounted on positively-charged glass slides and was subject to Nissl staining with cresyl violet as described in Paul et al. (43). Electrode lesions were photographed using a 5X brightfield microscope and digital camera (ThermoFisher, EVOS XL Core). Rats with inaccurate targeting were eliminated from the study.

## ACKNOWLEDGMENTS

This work was supported by grants from the National Institute of Mental Health (MH099085 to M.K.), the National Institute on Drug Abuse (DA043461 to M.K.), and the National Institute on Aging (AG049090 to A.W.). Select images were created with Biorender.com. We thank K. O’Reilly and A.A. Fenton for training KJS in electrode construction and rodent surgery. We thank A. Adhikari and M. van der Meer for sharing MATLAB scripts. We thank S. Mooer and T. Peterson for technical support.

