## Supplemental Figure 1 for "Sex differences and effects of estrous stage on hippocampal-prefrontal theta communications"

**SUPPORTING INFORMATION (SI)**

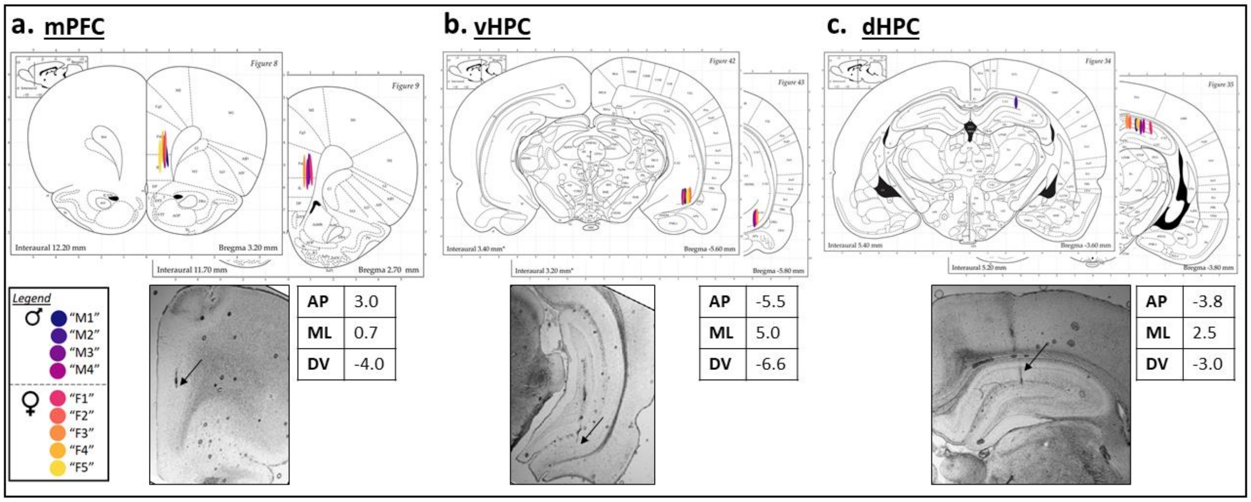


<u>**Figure S1.**</u> **Coordinate verification histology. (a)** Medial prefrontal cortex (mPFC). Four-electrode bundles with a total vertical spread of 3mm were implanted targeting mPFC-Infralimbic (IL), such that the two dorsal-most channels targeted mPFC-Prelimbic (PrL). **(b)** Ventral hippocampus (vHPC). Six-electrode bundles with a total vertical spread of 1mm were implanted targeting vHPC CA1. **(c)** Dorsal hippocampus (dHPC). Six-electrode bundles with a total vertical spread of 1mm were implanted targeting dHPC CA1. Male and female implant locations are matched. Tables list stereotaxic coordinates (AP: anterior-posterior; ML: medial-lateral; DV: dorsal-ventral). ***Below:*** Example photographs of electrode lesion sites, 5X objective. Black arrows indicate tissue lesions. Brain atlas images are adapted from Paxinos and Watson (38).
